# XSTREME: Comprehensive motif analysis of biological sequence datasets

**DOI:** 10.1101/2021.09.02.458722

**Authors:** Charles E. Grant, Timothy L. Bailey

## Abstract

XSTREME is a web-based tool for performing comprehensive motif discovery and analysis in DNA, RNA or protein sequences, as well as in sequences in user-defined alphabets. It is designed for both very large and very small datasets. XSTREME is similar to the MEME-ChIP tool, but expands upon its capabilities in several ways. Like MEME-ChIP, XSTREME performs two types of *de novo* motif discovery, and also performs motif enrichment analysis of the input sequences using databases of known motifs. Unlike MEME-ChIP, which ranks motifs based on their enrichment in the *centers* of the input sequences, XSTREME uses enrichment *anywhere* in the sequences for this purpose. Consequently, XSTREME is more appropriate for motif-based analysis of sequences regardless of how the motifs are distributed within the sequences. XSTREME uses the MEME and STREME algorithms for motif discovery, and the recently developed SEA algorithm for motif enrichment analysis. The interactive HTML output produced by XSTREME includes highly accurate motif significance estimates, plots of the positional distribution of each motif, and histograms of the number of motif matches in each sequences. XSTREME is easy to use via its web server at https://meme-suite.org, and is fully integrated with the widely-used MEME Suite of sequence analysis tools, which can be freely downloaded at the same web site for non-commercial use.

## 1 Introduction

Short, approximate sequence patterns (motifs) are known to encode functional biological signals in DNA, RNA and protein sequences. In genomic DNA, motifs capture the preferred binding sites of transcription factors (TFs) and promoter elements. In RNA motifs describe the binding preferences of RNA-binding proteins (RBPs). Motifs can also represent many protein features such as the targets of enzymes.

XSTREME is a web-based tool for comprehensive motif-based sequence analysis of DNA, RNA or protein data sets. It provides computationally efficient algorithms for discovering and analyzing the sequence motifs. Given a set of biological sequences, XSTREME first executes two different motif discovery algorithms: MEME (multiple EM for motif elicitation) [2], and STREME (Sensitive, Thorough, Rapid Enriched Motif Elicitation) [1] to discover novel sequence motifs. Next it then uses a motif enrichment analysis algorithm, SEA (Simple Enrichment Analysis) [3] to detect enrichment of previously characterized functional motifs and to rank the enrichment of discovered and known motifs on the same scale. Finally, to ease interpretation of the results, XSTREME applies a clustering algorithm to group the discovered and enriched motifs by similarity to each other. XSTREME returns its results as an interactive HTML document that provides visual representations of each motif and its locations within the input sequences, as well as estimates of the statistical significance of its enrichment. All results are presented in groups sorted according to statistical significance, and the HTML document provides clickable links to all details of the individual analyses. The XSTREME web-server provides numerous databases of known motifs for use in the motif enrichment analysis step, including motifs associated with given TFs (e.g, the JASPAR database [5]), motifs bound by RBPs (e.g., the CISBP-RNA database [9]), and protein motifs (ELM database [7]).

The XSTREME algorithm is similar to the existing MEME-ChIP [8], but XSTREME is applicable to a wider range of motif analysis problems. The primary distinction is that XSTREME makes no assumptions about the positional distribution of motif instances in the input sequences. XSTREME ranks motifs by how enriched they are in a primary set of sequences compared with a control set. (XSTREME will create a control set by shuffling the primary sequences if no control set is provided.) In contrast, MEME-ChIP ranks motifs by how enriched they are in the central regions of the input sequences. When it is believed (or known) that motifs may not be concentrated centrally in the sequences, XSTREME will provide more useful results than MEME-ChIP. Types of datasets where XSTREME is more appropriate than MEME-ChIP include sets of promoters, sets of accessible chromatin regions from ATAC-seq [4] and Cut&Run datasets using TF antibodies [6]. In each of these cases the assumption that motif motif sites will be near the centers of the input sequences does not always hold.

## 2 Results

Here, we illustrate using XSTREME for motif analysis of a Cut&Run dataset for the transcription factor Gata1 in erythroid precursor cells [11]. Cut&Run is an antibody-targeted cleavage method for identifying protein binding to chromatin from as few as 1000 cells [10]. In some experiments, however, Cut&Run results in a majority of bound sequences where the TF was near one end. With such datasets, XSTREME is a better fit than MEME-ChIP, which assumes that motifs tend to concentrate around the midpoint of the input sequences.

Fig. 1 shows the top motif cluster reported by XSTREME on the sequences with lengths from 400bp to 600bp specified in the Zhu *et al.* 2019 [11] dataset contained in file GSM4043375 GATA1 D7 S11 peaks.narrowPeak.gz (GEO accession number GSM4043375). The bi-modal distribution of the Gata1 binding sites is clearly apparent in the XSTREME output. The output shows that XSTREME has discovered a close match to the known Gata1 motif (the STREME motif 1-SAGATAAGR), and has associated that motif with the known Gata1 motif (MA0035.3) from the JASPAR database of motifs [5]. XSTREME also clusters both discovered and known motifs, and the known motif for the GATA1-TAL1 is shown aligned with the STREME motif. The figure also illustrates how XSTREME reports a histogram of the predicted number of motif matches per input sequence with at least one match. In this example, approximately 25% of the input sequences with one or more sites have multiple sites.

**Figure 1:**
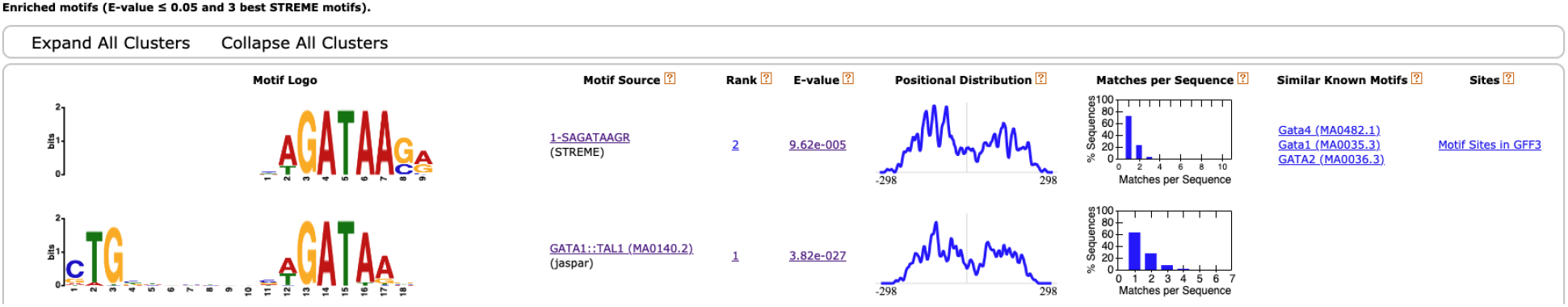
Screenshot of (a portion of) the HTML output of XSTREME run on Gata1 Cut&Run in erythroid precursor cell data.

